# iPSC-Derived Hepatocytes from Patients with MASLD Exhibit Early Mitochondrial Dysfunction

**DOI:** 10.1101/2025.08.27.672733

**Authors:** Dounia Le Guillou, Kevin Siao, Chris L. Her, Caroline C. Duwaerts, Jacquelyn J. Maher

## Abstract

**Background & Aims:** A hallmark of metabolic dysfunction-associated steatotic liver disease (MASLD) is a decline in the ability of hepatocyte mitochondria to adapt to excess lipid. This leads to the production of reactive oxygen species (ROS) and the instigation of a vicious cycle of further mitochondrial damage and cellular dysfunction that promotes disease progression. In this study, we investigated whether induced pluripotent stem cells (iPSCs) from MASLD patients exhibit features of mitochondrial dysfunction when differentiated to hepatocyte-like cells (iPSC-Heps).

**Methods:** iPSCs from 10 MASLD patients and 10 healthy control subjects genotyped for the I148M variant of PNPLA3 were differentiated to iPSC-Heps. Mitochondrial mass and function were assessed under basal culture conditions and following short-term exposure to exogenous palmitate. Outcomes included gene expression, mitochondrial oxygen consumption, ROS production and cellular energy status.

**Results:** iPSC-Heps from MASLD patients spontaneously accrued more lipid than control iPSC-Heps. Mitochondrial content was similar in MASLD and control iPSC-Heps, but MASLD iPSC-Heps displayed significant differences in mitochondrial function including a decrease in oxygen consumption rate when challenged with palmitate. Antioxidant gene expression was increased at baseline in MASLD vs. control iPSC-Heps, and MASLD iPSC-Heps produced more ROS and less ATP than controls after palmitate treatment. Differences persisted even when controlling for PNPLA3 genotype.

**Conclusions:** iPSC-Heps from MASLD patients exhibit mitochondrial alterations characteristic of their diseased origin. The degree of mitochondrial dysfunction seen in MASLD iPSC-Heps is reminiscent of that described clinically in early MASLD, prior to progression to steatohepatitis. Mitochondrial alterations in MASLD iPSC-Heps occur independently of PNPLA3 genotype.

## INTRODUCTION

Metabolic dysfunction-associated steatotic liver disease (MASLD) is one of the main causes of chronic liver disease worldwide.^1,2^ MASLD encompasses a spectrum of severity ranging from simple hepatic steatosis (MASL) to more severe steatohepatitis (MASH), the latter characterized by liver injury, inflammation and fibrosis.^3^ Although the exact mechanisms responsible for the onset of MASLD and the transition from MASL to MASH are not fully understood, it is known that excess lipid accumulation, oxidative stress and mitochondrial dysfunction are all important features of the disease.^4^

Mitochondria play a crucial role in fatty acid oxidation (FAO) and oxidative phosphorylation (OXPHOS), which are essential for energy production and cellular homeostasis. These processes are often altered in MASLD, leading to a disruption in cellular metabolism and the induction of cellular stress.^4-6^ Lipid accumulation in hepatocytes triggers mitochondrial dysfunction in MASLD by overloading mitochondrial FAO capacity.^4,7-10^ Indeed, mitochondrial dysfunction in hepatocytes is increasingly recognized as a key factor in disease progression from hepatic steatosis to MASH and fibrosis.^4,11,12^

Studies in humans have demonstrated that in the early stages of MASLD, hepatocyte mitochondria are able to adapt to lipid accumulation by raising their oxidative capacity^10-12^ and activating antioxidant defenses to efficiently scavenge reactive oxygen species (ROS) generated as by-products of OXPHOS.^13-15^ As MASL transitions toward MASH this mitochondrial adaptation deteriorates, leading to an inability to efficiently regulate the oxidative stress induced by chronic lipid overload. Excess mitochondrial ROS produced in this setting damage cellular DNA, protein and lipids, leading to lipid peroxidation and the activation of apoptotic and inflammatory signaling pathways.^16-21^ Moreover, excess ROS can impair mitochondrial biogenesis, which leads to a reduction of functioning mitochondrial mass.^11,12,22,23^ These observations underscore the importance of mitochondrial ROS as crucial drivers of MASLD/MASH and implicate ROS and mitochondrial dysfunction as pivotal components of a vicious cycle that promotes disease progression.^24,25^

The ability to generate induced pluripotent stem cells (iPSCs) from patients with liver disease and differentiate them to hepatocytes (iPSC-Heps) has enabled the interrogation of liver disease mechanisms without the need for invasive tissue procurement.^26-33^ Our group has utilized this approach to model human MASLD, using iPSCs reprogrammed from a cohort of patients with biopsy-proven MASLD – many of whom have MASH and liver fibrosis. We recently reported that iPSC-Heps from our MASLD subjects exhibit spontaneous steatosis in cell culture, whereas iPSC-Heps from healthy subjects do not.^34^ The objective of the current study was to extend this observation by determining whether MASLD iPSC-Heps also exhibit differences in mitochondrial profile compared to control iPSC-Heps. We find that MASLD iPSC-Heps display mitochondrial characteristics suggestive of a transition from MASL toward MASH evidenced by normal to increased respiratory capacity at baseline, but failure to boost respiration further in response to a lipid challenge in conjunction with increased ROS production. A factor likely influencing this disease-specific behavior is inheritance of rs738409, a single nucleotide polymorphism in *PNPLA3* (C>G) leading to an I148M substitution in adiponutrin that poses a significant MASLD risk.^35,36^ However, we could not attribute our results specifically to PNPLA3 genotype, as an examination of iPSC-Heps from all subjects who were homozygous for the *PNPLA3* GG variant did not reveal similar mitochondrial phenotypes. Instead, *PNPLA3* GG homozygotes exhibited distinct phenotypes corresponding to their original disease classification as control or MASLD, independent of their genetics.

## METHODS

### Generation of iPSCs and iPSC-Heps

Peripheral blood mononuclear cells were used to generate iPSCs from all subjects. iPSCs were reprogrammed by Cellular Dynamics International, Inc. (Novato, CA) using a virus-free, episomal reprogramming protocol as described by Yu and colleagues.^37^ The following cell lines from the CIRM iPSC Repository at Fujifilm were used in the study: CW10001, CW10002, CW10022, CW10030, CW10037, CW10077, CW10131, CW10149, CW10165, CW10166, CW10172, CW10173, CW10178, CW10182, CW10190, CW10192, CW10194, CW10201, CW10206 and CW10208. Subjects were genotyped for the I148M variant of *PNPLA3* (rs738409 C>G). iPSCs passed quality control testing for pluripotency and loss of reprogramming factors. iPSC-Heps were differentiated from iPSCs on multi-well plates coated with Matrigel® (Corning Life Sciences, Bedford, MA) using a modification of the Duncan protocol^38^ developed by Dr. A.N. Mattis.^39^ All experiments were performed at day 21 of differentiation. For Seahorse experiments, iPSCs were differentiated in Matrigel®-coated culture dishes until day 10, then detached with Accutase (Thermo Fisher Scientific, Waltham, MA) and replated at the same density in Seahorse XF24 V7 PS Cell Culture Microplates (Agilent, Santa Clara CA)

### Neutral Lipid Stain

iPSC-Heps were washed with warm phosphate-buffered saline (PBS), fixed with 4% paraformaldehyde for 20 min at room temperature and then washed three times with cold PBS. After fixation, cells were incubated with 1 μg/mL BODIPY 493/503 in PBS for 1 h at room temperature and then washed once with PBS. Nuclei were counterstained with 10 μg/mL Hoechst 33342 dye. Cells were imaged using a BioTek Cytation 5 imager.

### Cellular Triglyceride Measurement

Triglyceride levels in iPSC-Heps were measured using the Triglyceride Assay Kit from BioVision (Milpitas, CA) according to the manufacturer’s instructions. Briefly, iPSC-Heps were homogenized in 5% NP-40 and heated twice to 100 °C for 3 min followed by cooling to room temperature to ensure lipid solubilization. Samples were then centrifuged at 16,000 × g and supernatants were mixed with lipase for 20 min to release glycerol. Glycerol content was measured using a Synergy HT microplate reader (BioTek, Winooski, VT). Results were normalized to total cellular protein content (Pierce BCA Protein Assay, Thermo Fisher Scientific, Waltham, MA).

### Apolipoprotein B (APOB) Secretion

At the end of the differentiation protocol, culture supernatants from iPSC-Heps were collected and clarified by centrifugation. The concentration of APOB in the supernatants was quantified using the Human Apolipoprotein B ELISA Kit (Abcam, Cambridge, UK; Cat. #AB277393), according to the manufacturer’s instructions.

### RNA sequencing

RNA sequencing was performed on the iPSC-Heps in this study as part of a larger cohort.^34^ Reads were mapped to UCSC’s hg19 transcript set using Bowtie 2 version 2.1.0.^40^ Gene expression was quantified using RSEM’s (v1.2.15) expectation maximization algorithm.^41^ Quantified gene counts were normalized to counts per million, and then further normalized using the trimmed mean of M-values functionality in edgeR.^42^ Differential gene expression was analyzed by disease group, with batch effect addressed as a covariate in a generalized linear model.^42^ The RNA sequencing data set used in this study are deposited in GEO: GSE306392.

### Studies of mitochondrial mass and integrity

Mitochondrial mass was first assessed in live cells by staining with MitoTracker Green FM (Invitrogen, Waltham, MA) and quantitating fluorescence units per cell. Second, mitochondrial DNA (mtDNA) levels were measured by extracting total DNA from differentiated iPSC-Heps (QIAGEN DNeasy Blood & Tissue Kit, Redwood City, CA) and then performing qPCR for the cytochrome c oxidase subunit 1 (COX1) (PowerUp SYBR Green Master Mix, Thermo Fisher). Primers for this reaction *were 5′-TACGTTGTAGCCCACTTCCACT-3′* (forward) and *5′-AGTAACGTCGGGGCATTCCG-3′ (reverse)*. Values were normalized to ribosomal protein S6 (RPS6), quantitated using primers *5′-TGATGTCCGCCAGTATGTTG-3′* (forward) and *5′-TCTTGGTACGCTGCTTCTTC-3′ (reverse)*. Third, mitochondrial membrane integrity was assessed by measuring citrate synthase activity in whole cell extracts (Citrate Synthase Activity Assay Kit, Millipore Sigma, Burlington, MA). Enzyme activity was normalized to cellular protein content. To assess mitochondrial membrane potential after fatty acid exposure, iPSC-Heps were first treated for 2 h with BSA (control) or palmitate complexed to BSA (350 μM, 700 μM). Following treatment, the cells were washed with HBSS and then incubated in HBSS containing Hoechst 33342 and either 500 nM MitoTracker Red FM (membrane potential-dependent) or 200 nM Green FM (membrane potential-independent) dye (Invitrogen) for 20 min at 37°C and 5% CO_2_ in the dark. The staining solution was then removed, and cells were washed three times with HBSS. iPSC-Heps were imaged immediately using a BioTek Cytation 5 imager. Fluorescence intensity per cell (nucleus) was measured using CellProfiler image analysis software.

### Oxygen Consumption Measurements

Mitochondrial oxygen consumption rates (OCR) in the presence of electron transport chain inhibitors and uncouplers (oligomycin, 1.5 μM; carbonyl cyanide-p-trifluoromethoxyphenylhydrazone (FCCP), 2 μM; rotenone, 0.5 μM; antimycin A, 0.5 μM) were measured in adherent iPSC-Heps using a Seahorse XFe24 Analyzer (Agilent, Santa Clara, CA).Two assays were performed: the Mito Stress Test and the Seahorse XF Palmitate Oxidation Stress Test (Agilent) according to the manufacturer’s instructions. For the Mito Stress Test, fully differentiated iPSC-Heps were pre-incubated in DMEM supplemented with either 10 mM glucose and 2 mM glutamine (*glucose condition*) or 10 mM galactose and 6 mM glutamine (*galactose condition*) for 3 h to induce a metabolic switch.^43^ OCR was then assessed under the same substrate conditions plus 1mM pyruvate. For the Palmitate Oxidation Stress Test, iPSC-Heps were pre-incubated overnight in a substrate-limited medium containing 0.5 mM glucose, 0.5 mM L-carnitine and 1 mM GlutaMAX to promote fatty acid oxidation (FAO). Endogenous FAO was then assessed in the Seahorse Analyzer by comparing OCR in the presence and absence of etomoxir, an inhibitor of carnitine palmitoyltransferase-1. Exogenous FAO was assessed in parallel wells of iPSC-Heps by treating the cells with or without palmitate immediately prior to Seahorse analysis. Endogenous and exogenous FAO were calculated as the difference in OCR between cells treated with or without etomoxir and with or without palmitate, respectively. OCR values were normalized to the number of cells in each well, assessed by counting stained nuclei (Hoechst 33342, Sigma-Aldrich) with CellProfiler image analysis software.

### Assessment of Mitochondrial ROS Generation

Mitochondrial ROS generation in response to fatty acid treatment was assessed using MitoSOX Red dye, which detects mitochondrial superoxide. iPSC-Heps were treated for 2 h with either BSA (control) or palmitate complexed to BSA (350 μM or 700 μM). They were then washed with warm HBSS and incubated for 20 min with 3 μM MitoSOX Red at 37°C and 5% CO_2_ in the dark. Cells were gently washed once with warm HBSS buffer and fluorescence intensity measured using well-scanning spectrofluorimetry on a Synergy HT microplate reader at excitation/emission wavelengths of 520/590 nm.

### Cellular energy status

ADP/ATP ratios in iPSC-Heps were measured using the ADP/ATP Ratio Assay Kit (Sigma-Aldrich) according to the manufacturer’s instructions. Briefly, iPSC-Heps treated for 2 h with BSA or palmitate were lysed using the ATP reagent mix and luminescence was recorded to quantify ATP levels. After 10 minutes, ADP was converted to ATP, and luminescence was measured again to assess total content. The ADP/ATP ratio was then calculated from these measurements.

### Statistical analyses

In all assays, outcome measures were assessed using iPSC-Heps from 6-10 MASLD and 6-10 control subjects. The number of subjects analyzed in each assay is indicated in the figure legends. Statistical comparisons between MASLD and control groups were made using the Student’s t-test; linear regression was assessed using the F-test. In some cases, 2-way ANOVA was used with Šídák’s post hoc test for multiple comparisons. *P* values <.05 were considered significant.

### Author review and approval

All authors had access to the primary study data, and all authors reviewed and approved the final version of the manuscript.

## RESULTS

The 20 iPSC lines used in this study were selected from a group of 37 iPSC lines representing 21 patients with biopsy-proven MASLD and 16 healthy control subjects whose transcriptomic profiles were previously published by our group.^34^ The 20 cell lines were chosen based on their clear distinction into MASLD (n = 10) and healthy (n = 10) groups based on gene signatures following differentiation to iPSC-Heps, as shown in **Figure S1**.^34^ Select demographic and clinical features of the 20 iPSC donors are shown in **Table 1**. For all experiments, iPSCs were differentiated to iPSC-Heps over 21 days.

**Table 1.** Subject Demographics.

| Sample ID | Age | Sex | Ethnicity | BMI | DM2 | PNPLA3 Genotype | St (0-3) | Bal (0-2) | Inf (0-3) | MAS (0-8) | Fib (0-4) | Pathologist Diagnosis |
| --- | --- | --- | --- | --- | --- | --- | --- | --- | --- | --- | --- | --- |
| Control 1 | 23 | F | SA | 19.2 | N | **GG** | -- | -- | -- | -- | -- | -- |
| Control 2 | 23 | F | EA | 22.3 | N | **GG** | -- | -- | -- | -- | -- | -- |
| Control 3 | 31 | F | HW | 25.4 | N | GC | -- | -- | -- | -- | -- | -- |
| Control 4 | 32 | F | HW | 26.3 | N | **GG** | -- | -- | -- | -- | -- | -- |
| Control 5 | 32 | M | SA | 22.4 | N | **GG** | -- | -- | -- | -- | -- | -- |
| Control 6 | 37 | F | AA | 24.4 | N | CC | -- | -- | -- | -- | -- | -- |
| Control 7 | 38 | M | Mixed | 21.2 | N | CC | -- | -- | -- | -- | -- | -- |
| Control 8 | 38 | M | NHPI | 26.6 | N | CC | -- | -- | -- | -- | -- | -- |
| Control 9 | 42 | M | HW | 27.5 | N | CG | -- | -- | -- | -- | -- | -- |
| Control 10 | 47 | M | NHW | 25.0 | N | **GG** | -- | -- | -- | -- | -- | -- |
| MASLD 1 | 30 | F | NHW | 32.7 | N | **GG** | 3 | 0 | 0 | 3 | 0 | MASL |
| MASLD 2 | 38 | M | SA | 22.9 | N | CG | 1 | 1 | 1 | 3 | 1 | MASH |
| MASLD 3 | 45 | F | HW | 37.9 | N | **GG** | 2 | 2 | 1 | 5 | 3 | MASH |
| MASLD 4 | 47 | F | HW | 37.2 | Y | **GG** | 1 | 2 | 1 | 4 | 3 | MASH |
| MASLD 5 | 48 | M | HW | 30.0 | N | **GG** | 2 | 1 | 1 | 4 | 3 | MASH |
| MASLD 6 | 49 | M | HW | 22.1 | Y | CG | 2 | 1 | 1 | 4 | 3 | MASH |
| MASLD 7 | 52 | M | HW | 29.9 | Y | **GG** | 2 | 2 | 1 | 5 | 4 | MASH |
| MASLD 8 | 54 | F | HW | 43.7 | Y | **GG** | 1 | 2 | 1 | 4 | 3 | MASH |
| MASLD 9 | 57 | F | HW | 58.1 | Y | **GG** | 3 | 2 | 3 | 8 | 4 | MASH |
| MASLD 10 | 71 | F | HW | 36.0 | pre | **GG** | 1 | 2 | 1 | 4 | 4 | MASH |

SA, South Asian; EA, East Asian; NHW, non-Hispanic White; NHPI, Native Hawaiian or other Pacific Islander; HW, Hispanic White; Bal, ballooning; DM2, type 2 diabetes; Fib, fibrosis; Inf, inflammation; MAS, MASLD Activity Score; St, steatosis; MASL, metabolic dysfunction-associated steatosis; MASH, metabolic dysfunction-associated steatohepatitis

### MASLD iPSC-Heps accumulate more lipid than control iPSC-Heps

When differentiated into iPSC-Heps, MASLD iPSCs displayed spontaneous steatosis, whereas control iPSCs did not (**Figure 1A**). Quantitation of cellular triglyceride (TG) confirmed that the TG content of MASLD iPSC-Heps was significantly higher than that of control iPSC-Heps (**Figure 1B**). To determine whether steatosis in MASLD iPSC-Heps coincides with suppression of lipid export, we measured APOB – an essential component of very-low-density lipoprotein (VLDL) particles – in the culture medium. APOB levels were significantly lower in medium from MASLD than control iPSC-Heps (**Figure 1C**). Consistent with this finding, we found that *APOB* gene expression was 2.9-fold lower in MASLD than control iPSC-Heps (**Figure 1D**).

**Figure 1.**
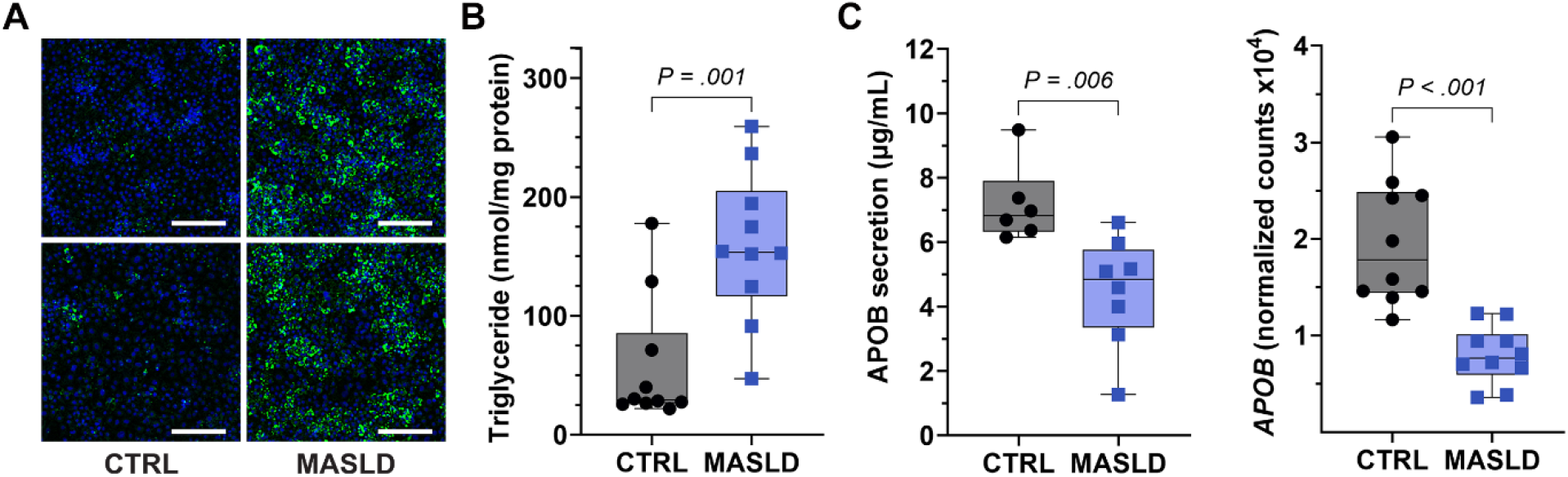
Lipid content and lipid secretion in MASLD vs. control (CTRL) iPSC-Heps. **(A)** Photomicrographs illustrate lipid droplets in 2 MASLD and 2 CTRL iPSC-Hep cultures at day 21 of differentiation, stained with BODIPY 493/503. Scale bar = 200 µm. **(B)** Graph depicts cellular triglyceride levels measured biochemically in MASLD (n = 10) and CTRL (n = 10) iPSC-Heps. **(C)** Graph illustrates APOB measured by ELISA in the cell culture supernatant from MASLD (n = 8) and control (n = 6) iPSC-Heps. **(D)** Graph depicts APOB mRNA levels in MASLD (n = 10) and CTRL (n = 10) iPSC-Heps. *P* values represent the results of unpaired Student’s t-tests.

### Mitochondrial mass is comparable in MASLD and control iPSC-Heps

To determine whether mitochondrial mass is different in MASLD vs. control iPSC-Heps, we stained cells with MitoTracker Green, a dye that identifies all mitochondria regardless of mitochondrial membrane potential. Fluorescence intensity was comparable in MASLD and control iPSC-Heps (**Figure 2A, B**). We also measured mitochondrial DNA (mtDNA) levels in MASLD and control iPSC-Heps and found no significant difference between the two groups (mtDNA/nDNA = 1.0 ± 0.2 vs. 0.8 ± 0.4 in control vs. MASLD, respectively). Citrate synthase, an enzyme localized within the mitochondrial matrix, was equally active in MASLD and control iPSC-Heps (**Figure 2C**), although expression of the citrate synthase gene (*CS*) was 16% higher in MASLD iPSC-Heps (**Figure 2D**). We did not detect differential expression of genes involved in mitochondrial biogenesis between MASLD and control iPSC-Heps (**Figure S2**).

**Figure 2.**
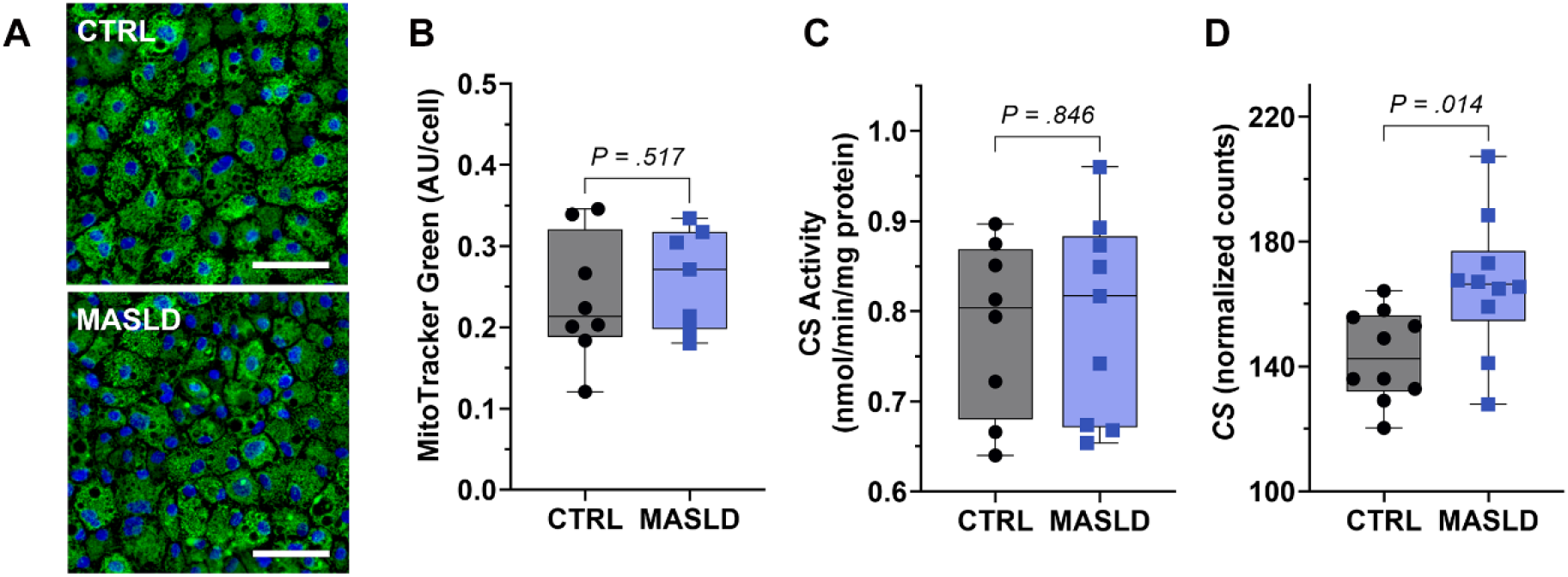
Mitochondrial content and integrity in MASLD vs. control (CTRL) iPSC-Heps. **(A)** Photomicrographs illustrate MASLD and CTRL iPSC-Heps stained with MitoTracker Green on day 21 of differentiation. Scale bar = 100 µm. **(B)** Graph depicts mitochondrial mass in MASLD (n=7) and CTRL (n=8) iPSC-Heps based on MitoTracker Green staining per cell, as outlined in Methods. **(C)** Graph illustrates citrate synthase (CS) activity in MASLD (n=8) and CTRL (n=9) iPSC-Heps. **(D)** Graph depicts CS gene expression in MASLD (n=10) and CTRL (n=10) iPSC-Heps. *P* values represent the results of unpaired Student’s t-tests.

### Mitochondrial respiration differs between MASLD and control iPSC-Heps

Having shown that mitochondrial mass is comparable in MASLD and control iPSC-Heps, we turned our attention to studies of cellular respiration. We first evaluated the expression of genes involved in oxidative phosphorylation (OXPHOS): among the 97 genes encoding components of complex I-V, we identified 19 that were differentially expressed between MASLD and control iPSC-Heps (**Figure 3A**). NDUF family genes, which encode the regulatory subunits of complex I of the mitochondrial respiratory chain (NADH dehydrogenase), showed mixed results, with some being upregulated and some downregulated. Genes encoding complex V, the ATP synthase, were not differentially expressed between the MASLD and control groups (not shown). Genes representing complexes II (succinate dehydrogenase) and IV (cytochrome c oxidase) were downregulated in MASLD vs. control iPSC-Heps. The latter is reminiscent of a recent finding in mice, in which liver-specific deletion of a complex IV subunit (Cox10) provoked hepatic steatosis and organ dysfunction.^44^

**Figure 3.**
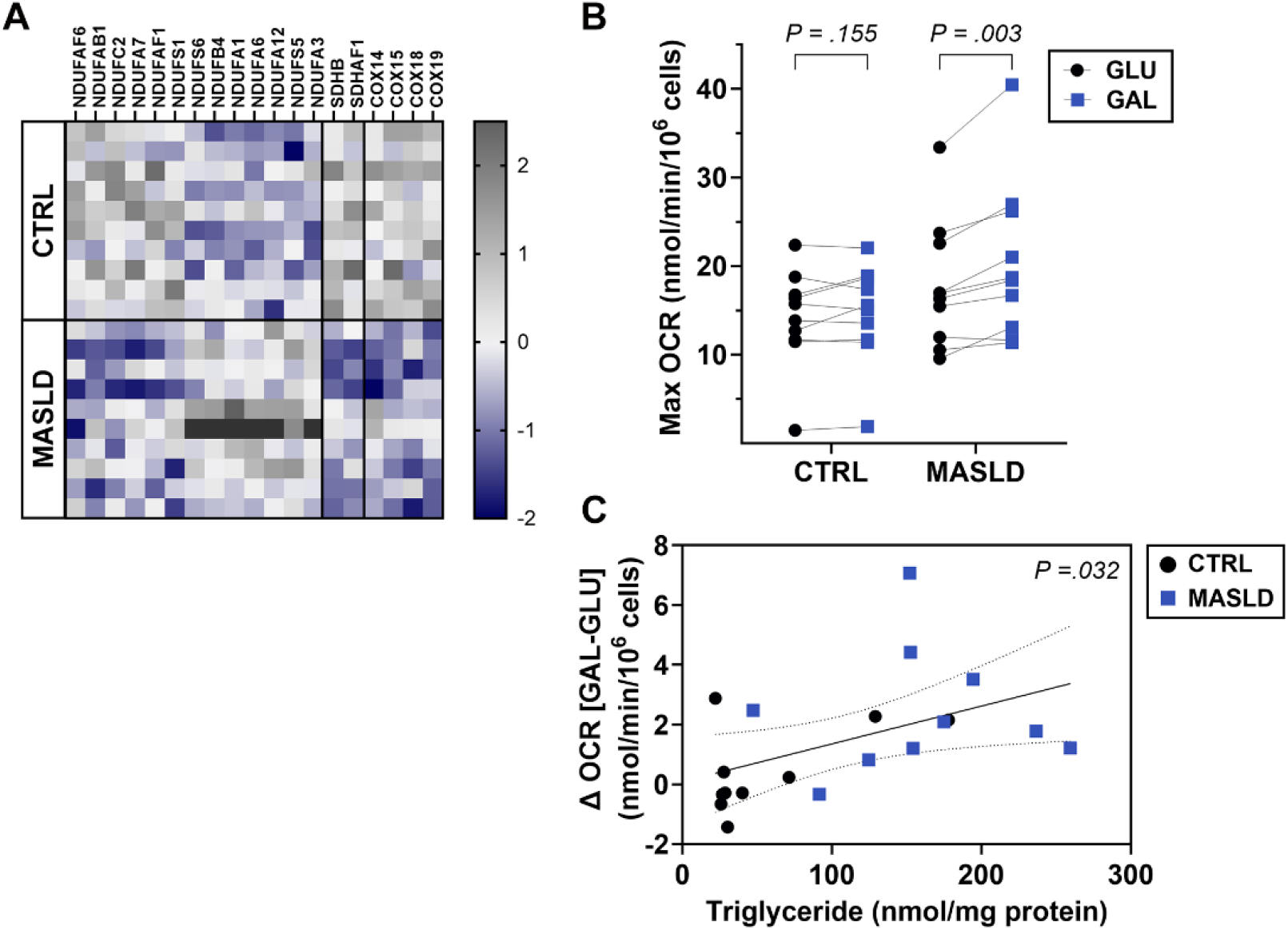
Measures of mitochondrial respiration in MASLD vs. control (CTRL) iPSC-Heps. **(A)** Heatmap illustrates genes involved in oxidative phosphorylation that are differentially expressed (*P* <.05) in MASLD (n = 10) vs. CTRL (n = 10) iPSC-Heps on day 21 of differentiation under standard culture conditions. **(B)** Graph depicts the maximal oxygen consumption rate (OCR) in MASLD (n=10) and CTRL (n=10) iPSC-Heps when pre-incubated in glucose (GLU) or galactose (GAL). *P* values represent the results of paired Student’s t-tests. **(C)** Graph demonstrates the correlation between the difference in OCR in galactose vs. glucose for each cell line (Δ OCR [GAL-GLU]) and cellular triglyceride content (n = 10 MASLD, n = 10 CTRL). The *P* value is the result of an F-test; dotted lines represent 95% confidence intervals.

We next sought to evaluate mitochondrial respiratory chain activity by measuring the mitochondrial oxygen consumption rate (OCR) in MASLD vs. control iPSC-Heps. We measured OCR in iPSC-Heps cultured under standard conditions with glucose (GLU) as well as in medium in which glucose was replaced by galactose (GAL). The latter condition limits glycolytic ATP production and promotes mitochondrial oxidative metabolism, thus allowing assessment of the maximal respiratory capacities of the cells without glycolytic support.^45^ Control iPSC-Heps showed no difference in maximal OCR (after addition of FCCP) whether under GLU or GAL conditions (**Figure 3B**). However, MASLD iPSC-Heps significantly increased their maximal OCR when cultured with GAL compared to GLU (**Figure 3B**), indicating that their mitochondrial respiratory capacity is unmasked when cells are forced to rely on oxidative metabolism and suggesting they are programmed differently metabolically than control iPSC-Heps. Interestingly, the change in OCR between GLU and GAL conditions (Δ OCR [GAL-GLU]) correlated positively with intracellular TG content (**Figure 3C**). This suggests that their increased lipid availability contributes to their enhanced mitochondrial substrate flux under oxidative conditions.

### Fatty acid oxidation in MASLD and control iPSC-Heps

To determine whether fatty acid oxidation (FAO) is dysregulated in MASLD vs. control iPSC-Heps, we first reviewed the relative expression of genes pertinent to this pathway. Under basal culture conditions, 19 genes involved in fatty acid degradation were significantly downregulated in MASLD iPSC-Heps (**Figure 4A**). Fifteen of the 19 genes encode enzymes located in the mitochondria; the remaining four encode enzymes located in peroxisomes (*ACOX1*) or cytosol (*ACAT2, ADH4, ADH5*). We then assessed FAO directly in iPSC-Heps by measuring OCR in the presence or absence of etomoxir, which prevents entry of fatty acids into the mitochondria and permits assessment of oxygen consumption attributable to endogenous fatty acids. Despite their lower expression of FAO-related genes, MASLD iPSC-Heps exhibited higher endogenous FAO-linked OCR than control iPSC-Heps (**Figure 4B**). This endogenous FAO was positively correlated with iPSC-Hep TG content (**Figure 4C**). On the contrary, we found that FAO-linked OCR when measured following the addition of exogenous palmitate was significantly lower in MASLD iPSC-Heps than controls (**Figure 4D**), and negatively correlated with cellular TG content (**Figure 4E**). This result suggests that MASLD iPSC-Heps, which already contain excess cytosolic lipid, are unable to accommodate an exogenous lipid challenge, signaling mitochondrial dysfunction.

**Figure 4.**
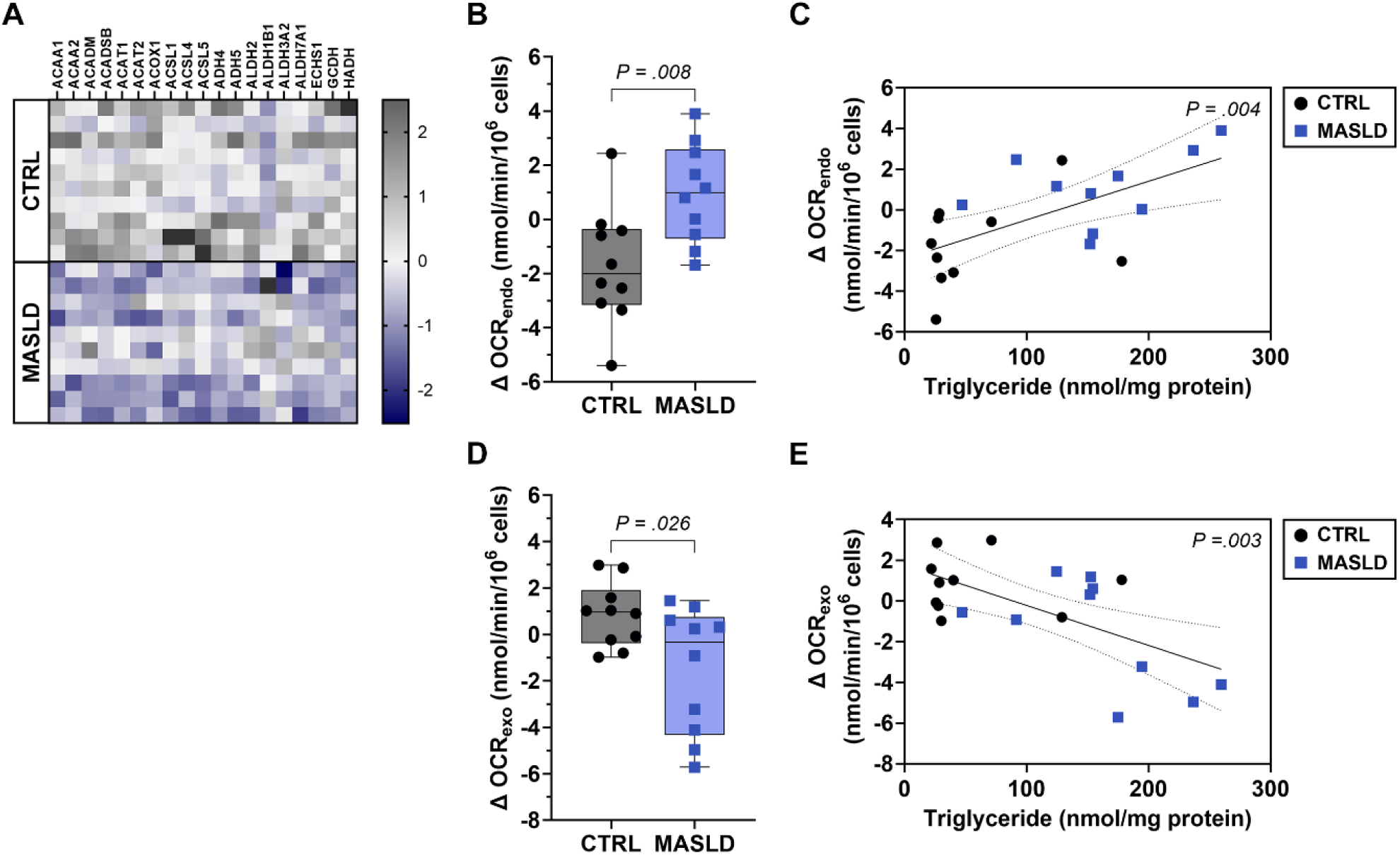
Fatty acid oxidation (FAO) in MASLD and control (CTRL) iPSC-Heps. **(A)** Heatmap illustrates genes involved in the oxidation of fatty acids that are differentially regulated (*P* <.05) in MASLD (n = 10) vs. CTRL (n = 10) iPSC-Heps on day 21 of differentiation under standard culture conditions. **(B)** Graph depicts the maximal endogenous FAO-linked OCR, measured as the difference in OCR measured with and without etomoxir (Δ OCR_endo_), in MASLD (n=10) and CTRL (n = 10) iPSC-Heps. **(C)** Graph demonstrates the correlation between the endogenous FAO-linked OCR shown in (B) and cellular triglyceride content (n = 10 MASLD, n = 10 CTRL). **(D)** Graph depicts the maximal exogenous FAO-linked OCR, calculated as the difference between conditions with and without palmitate supplementation (Δ OCR_exo_), in MASLD (n = 10) and control (n = 10) iPSC-Heps. **(E)** Graph demonstrates the correlation between the exogenous FAO-linked OCR shown in (D) and cellular triglyceride content (n = 10 MASLD, n = 10 CTRL). *P* values in (B) and (D) represent the results of paired Student’s t-tests. *P* values in (C) and (E) represent the results of F-tests; dotted lines represent 95% confidence intervals.

### Reactive oxygen species production in MASLD and control iPSC-Heps

Because mitochondrial dysfunction is typically associated with excess generation and accumulation of ROS, we evaluated ROS production and the related oxidative stress response in MASLD vs. control iPSC-Heps. A substantial number of genes involved in the response to oxidative stress, especially those that encode proteins responsible for ROS detoxification, were upregulated in MASLD vs. control iPSC-Heps, suggesting the presence of a demand for ROS neutralization in these cells (**Figure 5A**). Among the differentially expressed genes were those encoding superoxide dismutases (*SOD1, SOD3*) and glutathione peroxidases (*GPX2, GPX7*), as well as *KEAP1* (Kelch-like ECH-associated protein 1) and *NFE2L2* (Nuclear Factor Erythroid 2-Related Factor 2 or NRF2), which control the transcriptional activation of several antioxidant genes.^46^ Interestingly, the expression of most of those genes in MASLD and control iPSC-Heps was positively correlated with cellular TG content (**Figure 5B, Table S1**), which supports a link between lipid overload and oxidative stress responses. In contrast, Forkhead box O4 (*FOXO4*), a cell survival gene, was inversely correlated with iPSC-Hep TG content and oxidative stress genes (**Figure 5B, Table S1**). We then measured mitochondrial ROS production in MASLD and control iPSC-Heps after a 2-h treatment with 350 or 700 μM palmitate. Palmitate induced a significant increase in mitochondrial ROS production in MASLD iPSC-Heps, whereas it had no effect on control iPSC-Heps (**Figure 5C**).

**Figure 5.**
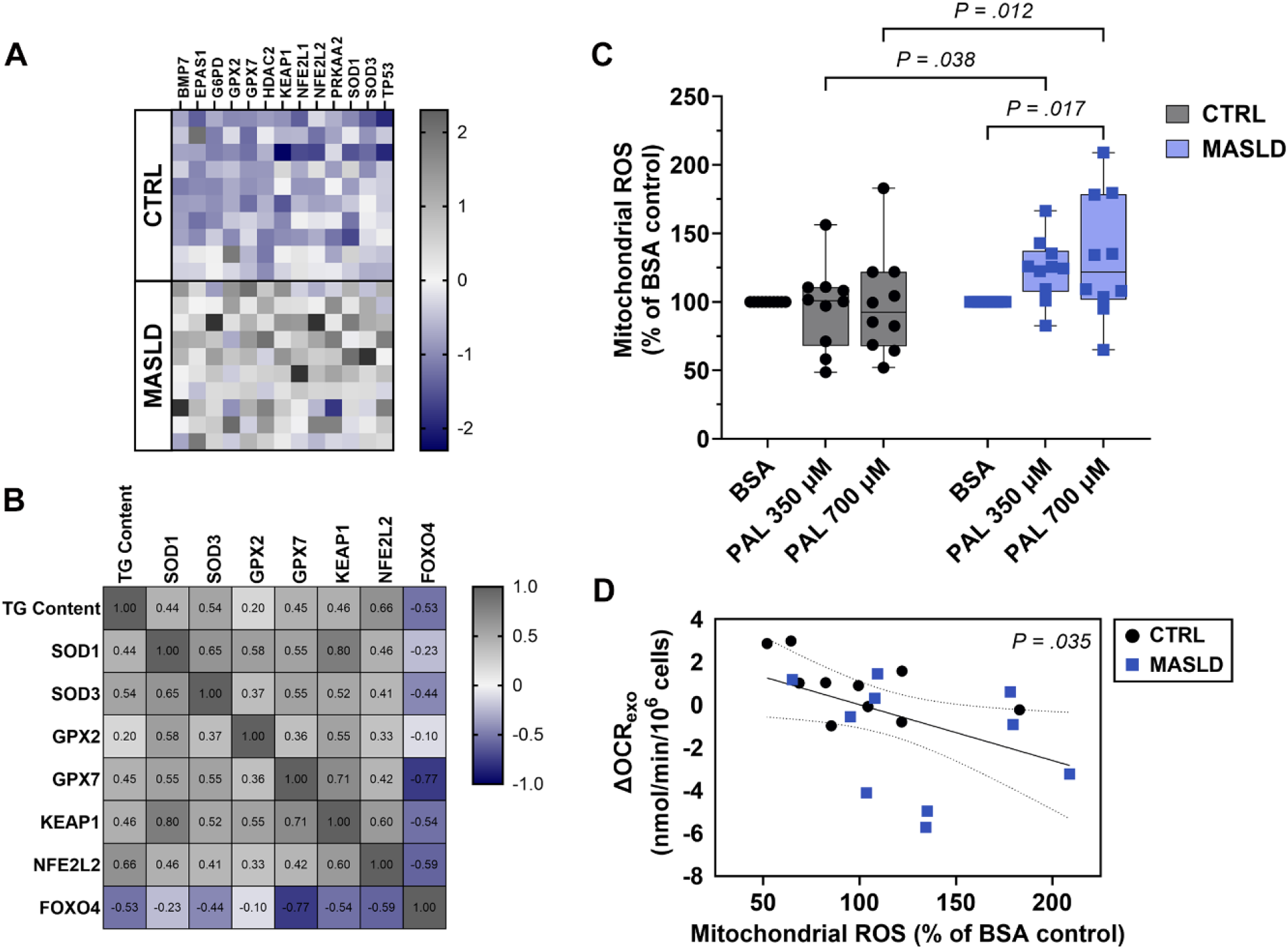
ROS production in MASLD vs. control (CTRL) iPSC-Heps. **(A)** Heatmap illustrates genes involved in the adaptive response to oxidative stress that are upregulated (*P* <.05) in MASLD (n = 10) vs. CTRL (n = 10) iPSC-Heps on day 21 of differentiation under standard culture conditions. **(B)** Heatmap illustrates the Pearson correlation between iPSC-Hep triglyceride (TG) content and oxidative stress-related gene expression. Pearson’s correlation coefficients and *P* values are indicated in Table S1. **(C)** Graph depicts mitochondrial ROS production in MASLD (n=10) and CTRL (n=10) iPSC-Heps treated with BSA or palmitate complexed to BSA (PAL). Two-way ANOVA identified a disease effect (*P* = 0.044); the illustrated *P* values are the result of Šídák’s post hoc tests. **(D)** Graph demonstrates the correlation between mitochondrial ROS production after 700 μM PAL treatment and the exogenous FAO-linked OCR shown in Figure 4D (n = 10 MASLD, n = 10 CTRL). The *P* value represents the result of an F-test; dotted lines represent 95% confidence intervals.

### ATP production in MASLD and control iPSC-Heps

We measured the ADP/ATP ratio in a subset of iPSC-Heps after 2 hours of treatment with BSA or palmitate to further evaluate mitochondrial energy production. In control iPSC-Heps, the ADP/ATP ratio tended to be lower in palmitate-treated cells (*P* = 0.08), indicating increased ATP generation in response to a lipid substrate (**Figure S3**). By contrast, there was no change in the ADP/ATP ratio in MASLD iPSC-Heps after palmitate treatment. In some MASLD lines, the ADP/ATP ratio was even increased, which suggests impaired coupling of fatty acid oxidation to ATP production or defective function of the electron transport chain in cells with lipid overload.

### Mitochondrial membrane potential in MASLD iPSC-Heps

The presence of altered mitochondrial OCR and increased ROS production in MASLD iPSC-Heps upon palmitate exposure led us to investigate whether mitochondrial membrane potential is altered in MASLD iPSC-Heps in response to a lipid challenge. We assessed the effect of acute palmitate treatment on mitochondrial membrane potential by measuring the ratio of MitoTracker Red (membrane potential-dependent) to Green (membrane potential-independent) fluorescence. In control iPSC-Heps, a 2-h exposure to palmitate did not induce any change in mitochondrial membrane potential compared to medium or BSA controls (**Figure 6 A, B**). In MASLD iPSC-Heps, however, the mitochondrial membrane potential increased after 2 h of palmitate treatment compared to the medium control (**Figure 6A, B**). This response is seemingly paradoxical, as depolarization would be expected under stress as suggested by the effect of palmitate on mitochondrial OCR (**Figure 4D**). However, hyperpolarization of mitochondria has been observed in cultured cells as the early phase of a biphasic response to pro-apoptotic agents.^47-49^

**Figure 6.**
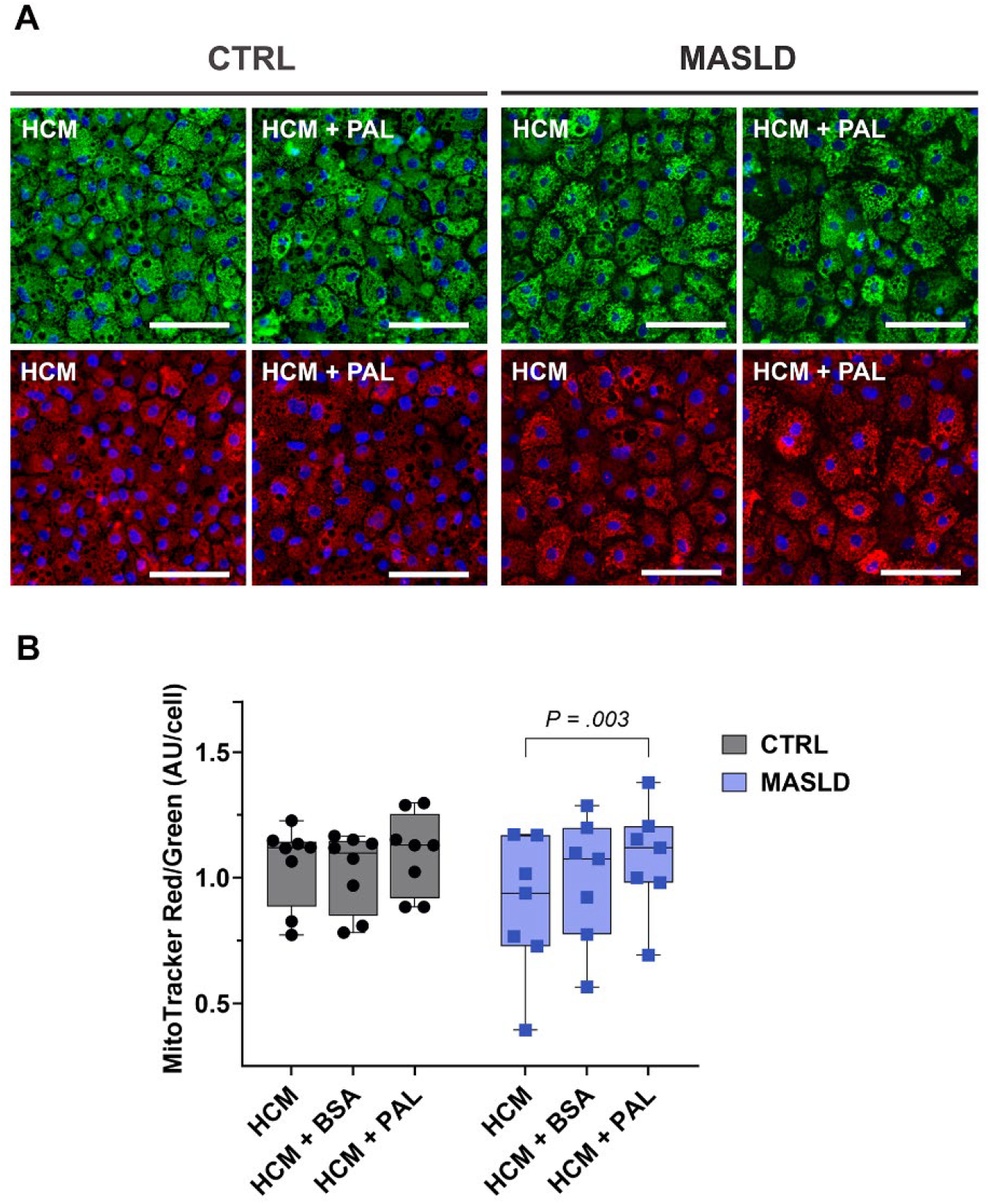
Effect of palmitate on mitochondrial membrane potential in MASLD vs. control (CTRL) iPSC-Heps. **(A)** Photomicrographs illustrate representative images of MASLD and CTRL iPSC-Heps stained with MitoTracker Green (membrane potential-independent) and MitoTracker Red (membrane potential-dependent) after a 2-h treatment with hepatocyte culture medium (HCM) or 700 µM palmitate complexed to BSA (HCM + PAL). Nuclei are stained with Hoechst 33342 (blue). Scale bar = 100 µm. **(B)** Quantitation of mitochondrial membrane potential, measured as the MitoTracker Red/Green ratio per cell in MASLD (n = 7) vs. CTRL (n = 8) iPSC-Heps, treated with HCM alone, HCM with BSA (HCM + BSA) or HCM supplemented with 700 µM palmitate complexed to BSA (HCM + PAL). Two-way repeated measures ANOVA identified a palmitate effect (*P* = 0.007); the illustrated *P* value represents the result of Šídák’s post hoc test.

### Lack of effect of the PNPLA3 variant rs738409 on APOB secretion or measures of mitochondrial function in iPSC-Heps

In the cohort of 20 subjects, 5 controls and 8 MASLD patients were homozygous for the PNPLA3 risk allele (G). To determine whether PNPLA3 genotype is a determinant of iPSC-Hep behavior regardless of the disease status of the donor, we performed a sub-analysis of experimental outcome measures in iPSC-Heps restricted to *PNPLA3* GG donors. The results indicate that APOB was still suppressed, endogenous FAO-related OCR was still increased and exogenous FAO-related OCR still decreased in MASLD vs. control iPSC-Heps, despite the *PNPLA3* GG genotype in all (**Figure 7A-D**). Similarly, MASLD iPSC-Heps from *PNPLA3* GG subjects produced more ROS in response to palmitate challenge than control subjects with the same genotype (**Figure 7E**). Notably, iPSC-Heps from MASLD subjects had significantly higher TG content than control iPSC-Heps despite a similar PNPLA3 genotype (**Figure 7F**).

**Figure 7.**
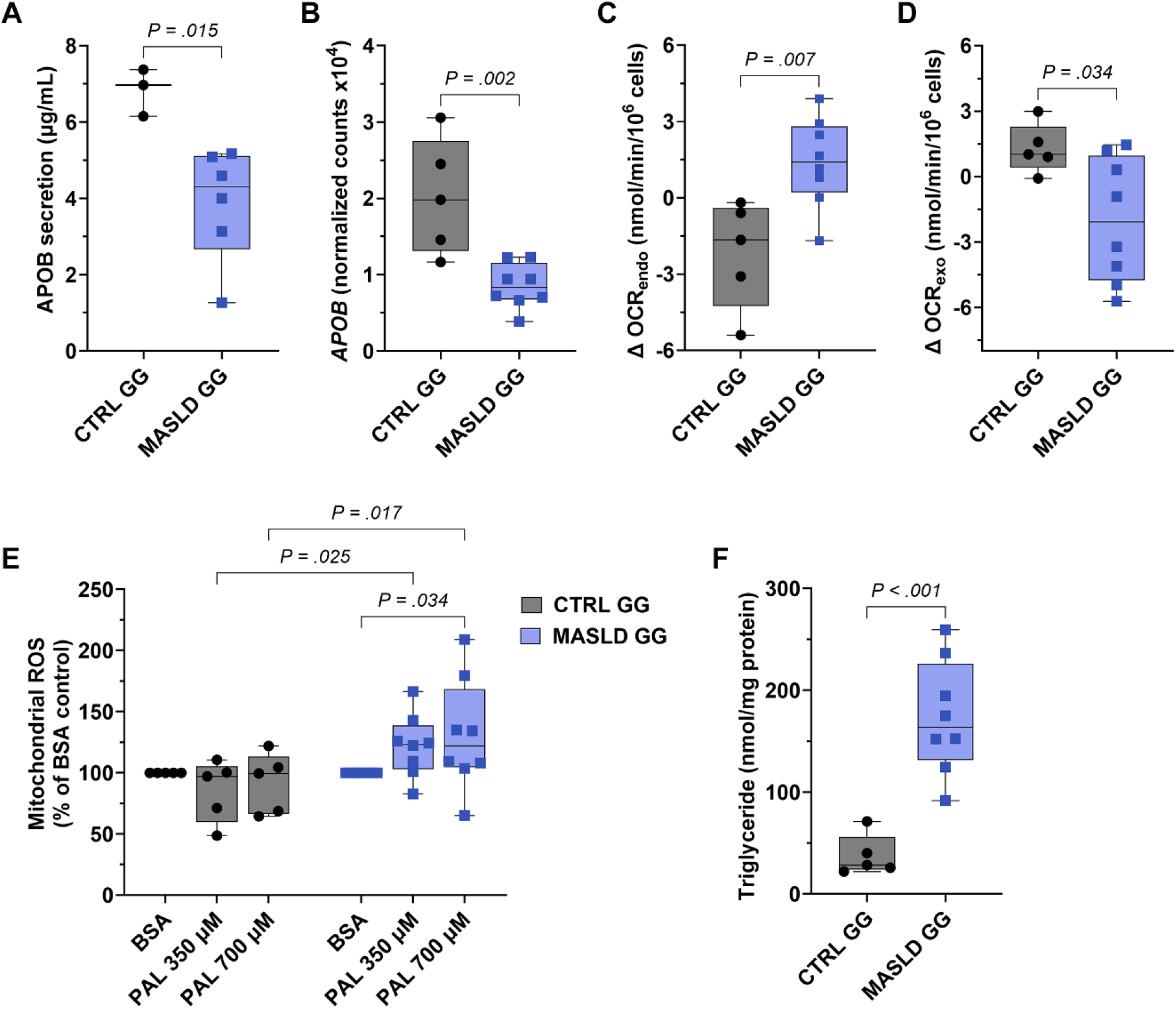
Sub-analyses of iPSC-Hep phenotype in MASLD vs. control iPSC-Heps from subjects homozygous for the I148M variant of PNPLA3 (*PNPLA3* GG). **(A, B)** Graphs depict APOB secretion and *APOB* expression as in Figure 1. **(C, D)** Graphs depict maximal endogenous FAO (Δ OCR_endo_) and exogenous FAO (Δ OCR_exo_) as in Figure 4. **(E)** Graph depicts ROS production in response to BSA or palmitate (PAL) as in Figure 5.

## DISCUSSION

Mitochondrial dysfunction plays a central role in the pathogenesis of MASLD and its progression to MASH.^4,50-52^ In this study we used iPSCs from 10 MASLD and 10 control subjects, selected specifically for their marked differences in transcriptomic profile after differentiation to iPSC-Heps,^34^ to look further into iPSC-Heps for disease-specific differences in mitochondrial function. Our experiments revealed significant differences between MASLD and control iPSC-Heps in mitochondrial oxygen consumption, fatty acid oxidation and oxidative stress responses. Indeed, MASLD iPSC-Heps displayed disturbances in mitochondrial function that align with those reported in human liver during the clinical progression of MASLD to MASH.^11^

Despite the use of identical culture conditions to differentiate all 20 iPSCs to iPSC-Heps, we showed that MASLD iPSC-Heps accrued more lipid during differentiation than control iPSC-Heps. In addition, MASLD iPSC-Heps in this cohort secreted significantly less APOB than control iPSC-Heps, which coincides with human studies demonstrating that circulating levels of APOB are reduced in a large proportion of patients with MASH.^53,54^ APOB secretion and triglyceride homeostasis were impaired in MASLD iPSC-Heps even in the absence of an exogenous metabolic challenge. This coincides with a theory proposed by Luukkonen and colleagues^55^ that suppression of VLDL synthesis in human MASLD occurs independently of substrate overload.

One central finding from the current study is that MASLD, but not control, iPSC-Heps increased their mitochondrial respiration when forced to use OXPHOS for ATP production. iPSC-Heps normally display active mitochondrial respiration;^33^ thus, they would not be expected to significantly change their oxygen consumption rate after only a brief exposure to galactose. The increase we observed in MASLD iPSC-Heps suggests that these cells have a low threshold for turning to endogenous fatty acids for energy production – a finding supported by the positive correlation between OCR and cellular lipid content. This reinforces the concept that mitochondrial respiration rises as an adaptive response to lipid accumulation in early MASLD.^10,11,56,57^ The increase in respiratory activity could represent an effort to control triglyceride accumulation and lipotoxicity; alternatively, it could reflect a decrease in mitochondrial respiratory efficiency necessitating an increase in respiratory activity to maintain adequate ATP production.^10,11^

Another finding of our study is the inability of MASLD iPSC-Heps to cope with an exogenous fatty acid load, as noted by their decrease in mitochondrial oxygen consumption rate and failure to generate additional ATP. This is in line with observations made in patients with MASH, who display a significant decrease in mitochondrial function compared to healthy subjects or patients with simple steatosis.^11,12^ When viewed together with the experiments documenting an increase in mitochondrial respiration in MASLD iPSC-Heps, our findings support a model in which steatosis causes a transient increase in mitochondrial function but reveals vulnerability to excessive or prolonged lipid accumulation. This mimics findings in rodents^58^ and humans^10,11^ that documented a transient increase in mitochondrial respiration during the development of MASLD that was then lost in MASH. In keeping with the mitochondrial dysfunction we observed in MASLD iPSC-Heps, our results revealed that these cells were experiencing oxidative stress based on elevated expression of genes involved in antioxidant defense. In rodents and humans, however, the progression from MASL to MASH typically coincides with a decrease, rather than increase, in antioxidant gene expression and activity.^11,59,60^ This discrepancy suggests that our MASLD iPSC-Heps mimic an early time point along the disease trajectory from MASL to MASH. Despite their higher levels of antioxidant defense genes under standard culture conditions, MASLD iPSC-Heps were not protected from ROS accumulation when challenged with palmitate, suggesting insufficient ROS detoxification capacity. Such ROS accumulation could fuel a vicious cycle culminating in more extensive oxidative damage and further mitochondrial dysfunction.^11,12,60,61^

Although MASLD iPSC-Heps showed signs of defective mitochondrial respiration (reduced OCR, **Figure 4D**) and increased ROS production (**Figure 5C**) when challenged with exogenous palmitate, we did not detect mitochondrial depolarization within a 2-hour treatment window (**Figure 6B**). Thus, our experiments revealed that mitochondrial stress appears very early in MASLD iPSC-Heps in response to palmitate, whereas mitochondrial depolarization may require more lengthy treatment. This is consistent with reports in the literature that palmitate can provoke mitochondrial depolarization in several types of cells including hepatocytes, but only after prolonged exposure, shortly before the onset of apoptosis.^62-67^ iPSC-Heps may also be able to maintain their mitochondrial integrity in the presence of palmitate due to their long-term adaptation to cell culture.

It is worth noting that we did not observe any clear change in mitochondrial mass between MASLD and control iPSC-Heps, nor did we find any alteration in factors involved in mitochondrial biogenesis (**Figure S2**). Reports in the literature regarding mitochondrial content in human MASH are mixed: some indicate content is increased compared to controls without liver disease,^11^ whereas others indicate no change^12,68^ and still others document an increase in citrate synthase activity but a decrease in mtDNA.^69^ Overall, our results support the findings reported in many human studies that mitochondrial function, but not content, is altered in human MASLD.

A key strength of this study is the fact that we were able to reproduce features of MASLD in patient-derived iPSC-Heps that heretofore could only be studied in liver tissue obtained from patients through invasive liver biopsy. Moreover, our experiments demonstrated that mitochondrial alterations are present consistently in iPSC-Heps from several unrelated MASLD donors, which underscores the utility of iPSC-based models to investigate the pathogenesis of the disease. Although the population investigated in this study was relatively small, it represents a disease cohort worthy of more complex disease modeling – such as the generation of heterotypic iPSC-based cell cultures that include MASLD hepatocytes, immune cells and hepatic stellate cells. A limitation of the study was our inability to definitively assess the impact of the PNPLA3 I148M gene polymorphism on the occurrence of mitochondrial alterations in MASLD iPSC-Heps, as has recently been proposed in humans.^70^ Our findings diverge from those of Luukkonen et al^70^ in that iPSC-Heps from our healthy control subjects expressing PNPLA3 I148M did not exhibit mitochondrial dysfunction despite their genetic risk. The reason for the discrepancy is unclear, but may be related to the lean body mass of our healthy PNPLA3 homozygotes.

In summary, our study demonstrates that iPSC-Heps from patients with biopsy-proven MASLD exhibit mitochondrial alterations representative of early MASLD in vivo. Several of these alterations were detectable in the absence of an exogenous metabolic challenge, which supports the notion that MASLD iPSC-Heps are intrinsically susceptible to metabolic stress. We were unable to attribute mitochondrial dysfunction in our MASLD iPSC-Heps to inheritance of the PNPLA3 I148M variant. In next steps, we hope to learn whether exposing MASLD iPSC-Heps to more prolonged substrate excesses similar to those experienced by MASLD patients unmasks further mitochondrial dysfunction. Future studies will also be directed toward uncovering whether MASLD-related genetic variants besides PNPLA3 I148M contribute to mitochondrial dysfunction in MASLD iPSC-Heps.

## Abbreviations

APOB: apolipoprotein B
ANOVA: analysis of variance
COX1: cytochrome oxidase subunit 1
CPT1: carnitine palmitoyl transferase 1
CTRL: control
FAO: fatty acid oxidation
GAL: galactose
GLU: glucose
iPSCs: induced pluripotent stem cells
iPSC-Heps: induced pluripotent stem cell-derived hepatocytes
HCM: hepatocyte culture medium
KEAP1: Kelch-like ECH-associated protein 1
KEGG: Kyoto Encyclopedia of Genes and Genomes
MASL: metabolic dysfunction-associated steatotic liver
MASLD: metabolic dysfunction-associated steatotic liver disease
MASH: metabolic dysfunction-associated steatohepatitis
mtDNA: mitochondrial DNA
NRF1: nuclear respiratory factor 1
OCR: oxygen consumption rate
OXPHOS: oxidative phosphorylation
PNPLA3: patatin-like phospholipase domain-containing protein 3
PPARGC1A: peroxisome proliferator-activated receptor gamma coactivator 1-alpha
ROS: reactive oxygen species
SNP: single nucleotide polymorphism
TFAM: transcription factor A, mitochondrial
TG: triglyceride
TM6SF2: transmembrane 6 superfamily member 2
VLDL: very-low-density lipoprotein

## Notes

**Grant support:** CIRM IT1-06563; NIH R21DK118380; NIH RC2DK136052; NIH P30DK026743

https://www.ncbi.nlm.nih.gov/geo/query/acc.cgi?acc=GSE306392

### Competing Interest Statement

The authors have declared no competing interest.

## REFERENCES

1. Younossi ZM. Non-alcoholic fatty liver disease - A global public health perspective. J Hepatol 2019;70:531–544.

2. Paik JM, Henry L, Younossi Y, et al. The burden of nonalcoholic fatty liver disease (NAFLD) is rapidly growing in every region of the world from 1990 to 2019. Hepatol Commun 2023;7.

3. Friedman SL, Neuschwander-Tetri BA, Rinella M, et al. Mechanisms of NAFLD development and therapeutic strategies. Nat Med 2018;24:908–922.

4. Fromenty B, Roden M. Mitochondrial alterations in fatty liver diseases. J Hepatol 2023;78:415–429.

5. Sharma S, Le Guillou D, Chen JY. Cellular stress in the pathogenesis of nonalcoholic steatohepatitis and liver fibrosis. Nat Rev Gastroenterol Hepatol 2023;20:662–678.

6. Shum M, Ngo J, Shirihai OS, et al. Mitochondrial oxidative function in NAFLD: Friend or foe? Mol Metab 2021;50:101134.

7. Serviddio G, Bellanti F, Tamborra R, et al. Alterations of hepatic ATP homeostasis and respiratory chain during development of non-alcoholic steatohepatitis in a rodent model. Eur J Clin Invest 2008;38:245–52.

8. Koves TR, Ussher JR, Noland RC, et al. Mitochondrial overload and incomplete fatty acid oxidation contribute to skeletal muscle insulin resistance. Cell Metab 2008;7:45–56.

9. Mantena SK, Vaughn DP, Andringa KK, et al. High fat diet induces dysregulation of hepatic oxygen gradients and mitochondrial function in vivo. Biochem J 2009;417:183–93.

10. Sunny NE, Parks EJ, Browning JD, et al. Excessive hepatic mitochondrial TCA cycle and gluconeogenesis in humans with nonalcoholic fatty liver disease. Cell Metab 2011;14:804–10.

11. Koliaki C, Szendroedi J, Kaul K, et al. Adaptation of hepatic mitochondrial function in humans with non-alcoholic fatty liver is lost in steatohepatitis. Cell Metab 2015;21:739–46.

12. Moore MP, Cunningham RP, Meers GM, et al. Compromised hepatic mitochondrial fatty acid oxidation and reduced markers of mitochondrial turnover in human NAFLD. Hepatology 2022;76:1452–1465.

13. Perlemuter G, Davit-Spraul A, Cosson C, et al. Increase in liver antioxidant enzyme activities in non-alcoholic fatty liver disease. Liver Int 2005;25:946–53.

14. Swiderska M, Maciejczyk M, Zalewska A, et al. Oxidative stress biomarkers in the serum and plasma of patients with non-alcoholic fatty liver disease (NAFLD). Can plasma AGE be a marker of NAFLD? Oxidative stress biomarkers in NAFLD patients. Free Radic Res 2019;53:841–850.

15. Monserrat-Mesquida M, Quetglas-Llabres M, Abbate M, et al. Oxidative Stress and Pro-Inflammatory Status in Patients with Non-Alcoholic Fatty Liver Disease. Antioxidants (Basel) 2020;9.

16. Ayala A, Munoz MF, Arguelles S. Lipid peroxidation: production, metabolism, and signaling mechanisms of malondialdehyde and 4-hydroxy-2-nonenal. Oxid Med Cell Longev 2014;2014:360438.

17. Berson A, De Beco V, Letteron P, et al. Steatohepatitis-inducing drugs cause mitochondrial dysfunction and lipid peroxidation in rat hepatocytes. Gastroenterology 1998;114:764–74.

18. Juan CA, Perez de la Lastra JM, Plou FJ, et al. The Chemistry of Reactive Oxygen Species (ROS) Revisited: Outlining Their Role in Biological Macromolecules (DNA, Lipids and Proteins) and Induced Pathologies. Int J Mol Sci 2021;22.

19. Win S, Than TA, L. BH, et al. Sab (Sh3bp5) dependence of JNK mediated inhibition of mitochondrial respiration in palmitic acid induced hepatocyte lipotoxicity. J Hepatol 2015;62:1367–74.

20. Satapati S, Kucejova B, Duarte JA, et al. Mitochondrial metabolism mediates oxidative stress and inflammation in fatty liver. J Clin Invest 2015;125:4447–62.

21. Bhogal RH, Curbishley SM, Weston CJ, et al. Reactive oxygen species mediate human hepatocyte injury during hypoxia/reoxygenation. Liver Transpl 2010;16:1303–13.

22. Aharoni-Simon M, Hann-Obercyger M, Pen S, et al. Fatty liver is associated with impaired activity of PPARgamma-coactivator 1alpha (PGC1alpha) and mitochondrial biogenesis in mice. Lab Invest 2011;91:1018–28.

23. Sheldon RD, Meers GM, Morris EM, et al. eNOS deletion impairs mitochondrial quality control and exacerbates Western diet-induced NASH. Am J Physiol Endocrinol Metab 2019;317:E605–E616.

24. Begriche K, Igoudjil A, Pessayre D, et al. Mitochondrial dysfunction in NASH: causes, consequences and possible means to prevent it. Mitochondrion 2006;6:1–28.

25. Simoes ICM, Fontes A, Pinton P, et al. Mitochondria in non-alcoholic fatty liver disease. Int J Biochem Cell Biol 2018;95:93–99.

26. Kimura M, Iguchi T, Iwasawa K, et al. En masse organoid phenotyping informs metabolic-associated genetic susceptibility to NASH. Cell 2022;185:4216–4232 e16.

27. Ouchi R, Togo S, Kimura M, et al. Modeling Steatohepatitis in Humans with Pluripotent Stem Cell-Derived Organoids. Cell Metab 2019;30:374–384 e6.

28. Takebe T, Sekine K, Kimura M, et al. Massive and Reproducible Production of Liver Buds Entirely from Human Pluripotent Stem Cells. Cell Rep 2017;21:2661–2670.

29. Collin de l’Hortet A, Takeishi K, Guzman-Lepe J, et al. Generation of Human Fatty Livers Using Custom-Engineered Induced Pluripotent Stem Cells with Modifiable SIRT1 Metabolism. Cell Metab 2019;30:385–401 e9.

30. Faccioli LAP, Sun Y, Animasahun O, et al. Human-induced pluripotent stem cell-based hepatic modeling of lipid metabolism-associated TM6SF2-E167K variant. Hepatology 2024.

31. Rezvani M, Vallier L, Guillot A. Modeling Nonalcoholic Fatty Liver Disease in the Dish Using Human-Specific Platforms: Strategies and Limitations. Cell Mol Gastroenterol Hepatol 2023;15:1135–1145.

32. Tilson SG, Morell CM, Lenaerts AS, et al. Modeling PNPLA3-Associated NAFLD Using Human-Induced Pluripotent Stem Cells. Hepatology 2021;74:2998–3017.

33. Jing R, Corbett JL, Cai J, et al. A Screen Using iPSC-Derived Hepatocytes Reveals NAD(+) as a Potential Treatment for mtDNA Depletion Syndrome. Cell Rep 2018;25:1469–1484 e5.

34. Duwaerts CC, Le Guillou D, Her CL, et al. Induced Pluripotent Stem Cell-derived Hepatocytes From Patients With Nonalcoholic Fatty Liver Disease Display a Disease-specific Gene Expression Profile. Gastroenterology 2021;160:2591–2594 e6.

35. Romeo S, Kozlitina J, Xing C, et al. Genetic variation in PNPLA3 confers susceptibility to nonalcoholic fatty liver disease. Nature genetics 2008;40:1461–5.

36. Kozlitina J, Sookoian S. Global Epidemiological Impact of PNPLA3 I148M on Liver Disease. Liver Int 2025;45:e16123.

37. Yu J, Chau KF, Vodyanik MA, et al. Efficient feeder-free episomal reprogramming with small molecules. PLoS One 2011;6:e17557.

38. Si-Tayeb K, Noto FK, Nagaoka M, et al. Highly efficient generation of human hepatocyte-like cells from induced pluripotent stem cells. Hepatology 2010;51:297–305.

39. Peaslee C, Esteva-Font C, Su T, et al. Doxycycline Significantly Enhances Induction of Induced Pluripotent Stem Cells to Endoderm by Enhancing Survival Through Protein Kinase B Phosphorylation. Hepatology 2021;74:2102–2117.

40. Langmead B, Salzberg SL. Fast gapped-read alignment with Bowtie 2. Nat Methods 2012;9:357–9.

41. Li B, Dewey CN. RSEM: accurate transcript quantification from RNA-Seq data with or without a reference genome. BMC Bioinformatics 2011;12:323.

42. Robinson MD, McCarthy DJ, Smyth GK. edgeR: a Bioconductor package for differential expression analysis of digital gene expression data. Bioinformatics 2010;26:139–40.

43. Kamalian L, Douglas O, Jolly CE, et al. Acute Metabolic Switch Assay Using Glucose/Galactose Medium in HepaRG Cells to Detect Mitochondrial Toxicity. Curr Protoc Toxicol 2019;80:e76.

44. Lesner NP, Wang X, Chen Z, et al. Differential requirements for mitochondrial electron transport chain components in the adult murine liver. Elife 2022;11.

45. Skolik RA, Solocinski J, Konkle ME, et al. Global changes to HepG2 cell metabolism in response to galactose treatment. Am J Physiol Cell Physiol 2021;320:C778–C793.

46. Nguyen T, Nioi P, Pickett CB. The Nrf2-antioxidant response element signaling pathway and its activation by oxidative stress. J Biol Chem 2009;284:13291–5.

47. Sanchez-Alcazar JA, Ault JG, Khodjakov A, et al. Increased mitochondrial cytochrome c levels and mitochondrial hyperpolarization precede camptothecin-induced apoptosis in Jurkat cells. Cell Death Differ 2000;7:1090–100.

48. Giovannini C, Matarrese P, Scazzocchio B, et al. Mitochondria hyperpolarization is an early event in oxidized low-density lipoprotein-induced apoptosis in Caco-2 intestinal cells. FEBS Lett 2002;523:200–6.

49. Cao J, Liu Y, Jia L, et al. Curcumin induces apoptosis through mitochondrial hyperpolarization and mtDNA damage in human hepatoma G2 cells. Free Radic Biol Med 2007;43:968–75.

50. Begriche K, Massart J, Robin MA, et al. Mitochondrial adaptations and dysfunctions in nonalcoholic fatty liver disease. Hepatology 2013;58:1497–507.

51. Loomba R, Friedman SL, Shulman GI. Mechanisms and disease consequences of nonalcoholic fatty liver disease. Cell 2021;184:2537–2564.

52. Ramanathan R, Ali AH, Ibdah JA. Mitochondrial Dysfunction Plays Central Role in Nonalcoholic Fatty Liver Disease. Int J Mol Sci 2022;23.

53. Charlton M, Sreekumar R, Rasmussen D, et al. Apolipoprotein synthesis in nonalcoholic steatohepatitis. Hepatology 2002;35:898–904.

54. Martinez-Arranz I, Bruzzone C, Noureddin M, et al. Metabolic subtypes of patients with NAFLD exhibit distinctive cardiovascular risk profiles. Hepatology 2022;76:1121–1134.

55. Luukkonen PK, Qadri S, Ahlholm N, et al. Distinct contributions of metabolic dysfunction and genetic risk factors in the pathogenesis of non-alcoholic fatty liver disease. J Hepatol 2022;76:526–535.

56. Jelenik T, Kaul K, Sequaris G, et al. Mechanisms of Insulin Resistance in Primary and Secondary Nonalcoholic Fatty Liver. Diabetes 2017;66:2241–2253.

57. Hyotylainen T, Jerby L, Petaja EM, et al. Genome-scale study reveals reduced metabolic adaptability in patients with non-alcoholic fatty liver disease. Nat Commun 2016;7:8994.

58. Satapati S, Sunny NE, Kucejova B, et al. Elevated TCA cycle function in the pathology of diet-induced hepatic insulin resistance and fatty liver. J Lipid Res 2012;53:1080–92.

59. Sreekumar R, Rosado B, Rasmussen D, et al. Hepatic gene expression in histologically progressive nonalcoholic steatohepatitis. Hepatology 2003;38:244–51.

60. Rector RS, Thyfault JP, Uptergrove GM, et al. Mitochondrial dysfunction precedes insulin resistance and hepatic steatosis and contributes to the natural history of non-alcoholic fatty liver disease in an obese rodent model. J Hepatol 2010;52:727–36.

61. Videla LA, Rodrigo R, Orellana M, et al. Oxidative stress-related parameters in the liver of non-alcoholic fatty liver disease patients. Clin Sci (Lond) 2004;106:261–8.

62. Sparagna GC, Hickson-Bick DL, Buja LM, et al. A metabolic role for mitochondria in palmitate-induced cardiac myocyte apoptosis. Am J Physiol Heart Circ Physiol 2000;279:H2124–32.

63. Egnatchik RA, Leamy AK, Jacobson DA, et al. ER calcium release promotes mitochondrial dysfunction and hepatic cell lipotoxicity in response to palmitate overload. Mol Metab 2014;3:544–53.

64. Moravcova A, Cervinkova Z, Kucera O, et al. The effect of oleic and palmitic acid on induction of steatosis and cytotoxicity on rat hepatocytes in primary culture. Physiol Res 2015;64:S627–36.

65. Sergi D, Luscombe-Marsh N, Naumovski N, et al. Palmitic Acid, but Not Lauric Acid, Induces Metabolic Inflammation, Mitochondrial Fragmentation, and a Drop in Mitochondrial Membrane Potential in Human Primary Myotubes. Front Nutr 2021;8:663838.

66. Schmitt LO, Blanco A, Lima SV, et al. Palmitate Compromises C6 Astrocytic Cell Viability and Mitochondrial Function. Metabolites 2024;14.

67. Ibrahim SH, Akazawa Y, Cazanave SC, et al. Glycogen synthase kinase-3 (GSK-3) inhibition attenuates hepatocyte lipoapoptosis. J Hepatol 2011;54:765–72.

68. Pedersen JS, Rygg MO, Chrois K, et al. Influence of NAFLD and bariatric surgery on hepatic and adipose tissue mitochondrial biogenesis and respiration. Nat Commun 2022;13:2931.

69. Gancheva S, Kahl S, Pesta D, et al. Impaired Hepatic Mitochondrial Capacity in Nonalcoholic Steatohepatitis Associated With Type 2 Diabetes. Diabetes Care 2022;45:928–937.

70. Luukkonen PK, Porthan K, Ahlholm N, et al. The PNPLA3 I148M variant increases ketogenesis and decreases hepatic de novo lipogenesis and mitochondrial function in humans. Cell Metab 2023;35:1887–1896 e5.

