## SUPPLEMENTAL FIGURES AND TABLES for "iPSC-Derived Hepatocytes from Patients with MASLD Exhibit Early Mitochondrial Dysfunction"

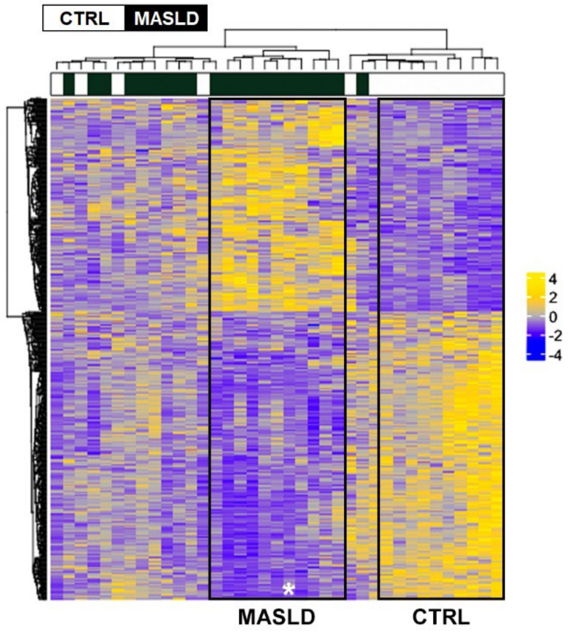


**Figure S1. Transcriptomic profiles of iPSC-Heps selected for the current study.** Heatmap illustrates the comparative gene expression of iPSC-Heps differentiated from 37 iPSC lines representing 21 MASLD patients and 16 healthy control subjects (adapted from reference 34). The framed boxes highlight the subjects selected for the current study, based upon their distinct transcriptomic profiles. Asterisk identifies one MASLD subject not included in the current analysis.

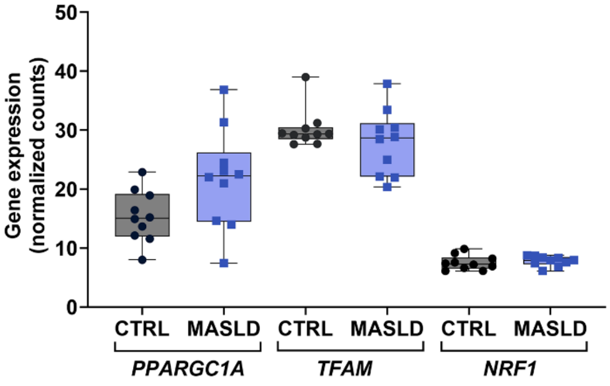


**Figure S2. Expression of genes involved in mitochondrial biogenesis in MASLD vs. control iPSC-Heps.** Graph depicts the expression of 3 genes involved in mitochondrial biogenesis, measured in MASLD (n=10) and CTRL (n=10) iPSC-Heps at day 21 of differentiation. No significant differences were observed. *PPARGC1A*, peroxisome proliferator-activated receptor gamma coactivator 1-alpha; *TFAM*, transcription factor A, mitochondrial; *NRF1*, nuclear respiratory factor 1.

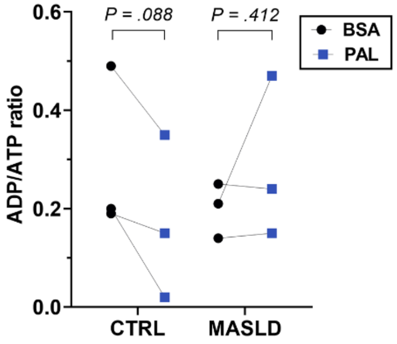


**Figure S3. Change in ADP/ATP ratio in MASLD vs. control iPSC-Heps in response to palmitate treatment.** Graph depicts the ADP/ATP ratio in MASLD (n = 3) and control (n = 3) iPSC-Heps following treatment with BSA or 700 μM palmitate (PAL). *P* values represent the results of paired t-tests.

**Table S1. Pearson's correlation between cellular triglyceride (cell TG) content and antioxidant gene expression.** Pearson's r values are shown for each pair. Entries in bold italic depict correlations with $P < .05$.

| | Cell TG | *SOD1* | *SOD3* | *GPX2* | *GPX7* | *KEAP1* | *NFE2L2* | *FOXO4* |
|---|---|---|---|---|---|---|---|---|
| **Cell TG** | 1.00 | 0.44 | ***0.54*** | 0.20 | ***0.45*** | ***0.46*** | ***0.66*** | ***-0.53*** |
| ***SOD1*** | 0.44 | 1.00 | ***0.65*** | ***0.58*** | ***0.55*** | ***0.80*** | ***0.46*** | -0.23 |
| ***SOD3*** | ***0.54*** | ***0.65*** | 1.00 | 0.37 | ***0.55*** | ***0.52*** | 0.41 | -0.44 |
| ***GPX2*** | 0.20 | ***0.58*** | 0.37 | 1.00 | 0.36 | ***0.55*** | 0.33 | -0.10 |
| ***GPX7*** | ***0.45*** | ***0.55*** | ***0.55*** | 0.36 | 1.00 | ***0.71*** | 0.42 | ***-0.77*** |
| ***KEAP1*** | ***0.46*** | ***0.80*** | ***0.52*** | ***0.55*** | ***0.71*** | 1.00 | ***0.60*** | ***-0.54*** |
| ***NFE2L2*** | ***0.66*** | ***0.46*** | 0.41 | 0.33 | 0.42 | ***0.60*** | 1.00 | ***-0.59*** |
| ***FOXO4*** | **-0.53** | -0.23 | -0.44 | -0.10 | **-0.77** | ***-0.54*** | **-0.59** | 1.00 |
